# Exploring Genomic Large Language Models: Bridging the Gap between Natural Language and Gene Sequences

**DOI:** 10.1101/2024.02.26.581496

**Authors:** Huaqing Liu, Shuxian Zhou, Peiyi Chen, Jiahui Liu, Ku-Geng Huo, Lanqing Han

**Author notes:** These authors contributed to equally the work (Huaqing Liu, Shuxian Zhou).

## Abstract

**Motivation:** With the rapid development of genomic sequencing technologies and accumulation of sequencing data, there is an increasing demand for analysis tools that are more user-friendly for non-programmer users. In support of this initiative, we developed an all-in-one tool called GenomicLLM that can understand simple grammar in the question input and perform different types of analyses and tasks accordingly.

**Reaults:** We trained the GenomicLLM model using three large open-access datasets, namely GenomicLLM_GRCh38, Genome Understanding Evaluation and GenomicBenchmarks, and developed a hybrid tokenization approach to allow better comprehension from mixed corpora that include sequence and non-sequence inputs. GenomicLLM can carry out a wider range of tasks. In the classification tasks that are also available in the state-of-the-art DNABERT-2 and HyenaDNA, GenomicLLM has comparable performance. Moreover, GenomicLLM can also carry out other regression and generation tasks that are not accomplishable by these tools. In summary, we demonstrated here a successful large language model with a mixture of gene sequences and natural language corpus that enables a wider range of applications.

**Availability and implementation:** Codes and data can be accessed at https://github.com/Huatsing-Lau/GenomicLLM and https://zenodo.org/records/10695802

## Introduction

Since the landmark completion of the initial human genome sequencing by the Human Genomic Project in 2003, there has been rapid accumulation of genomic, transcriptomic and epigenomic data, facilitated by the rapid developments in next-generation sequencing (NGS) technologies, and a surge in bioinformatic and artificial intelligence (AI)/machine learning (ML) tools for -omic data analysis.

These AI/ML tools allow different types of data analyses and predictions including promotor/enhancer prediction, splice site prediction, etc. However, many of these tools are either run in programming languages or have complicated interface that can be challenging to use. Moreover, most tools were designed to perform a single type or only a few types of tasks. There are also all-in-one platforms like R that allow users to perform a broad variety of tasks using different packages. However, it was designed for programmers who can operate in the R language and it would require users to download different packages for different types of tasks.

Large language models (LLM) like Generative Pre-trained Transformer (GPT) represent a significant advancement over traditional AI models due to their deep learning architecture and extensive pre-training on diverse datasets. Unlike conventional models that often require specific programming for each task, GPT models are designed to generalize across a wide range of tasks without task-specific training. One of the key advantages of GPT models is their ability to understand and generate human-like text, or natural language, making them highly versatile for applications like conversation, content creation, and language translation. The combination of generalizability and human-like text understanding makes LLMs a perfect tool to perform all-in-one -omic data analyses and predictions using simple human-like natural language questions.

There are tools like DNABERT, DNABERT-2 (Zhou, et al., 2023), DNAGPT, GeneGPT and HyenaDNA (Grešová, et al., 2023) that can understand nucleotide/amino acid sequence inputs and perform analysis to answer questions. However, these tools either require programming languages or have restricted input formats. Natural language models such as ChatGPT and LLaMA (Touvron, et al., 2023) can understand natural language input as questions and provide answers in a human-like style. However, they are not able to analyze nucleotide/amino acid sequence inputs. In the current paper, we developed an all-in-one tool called GenomicLLM that enables us to carry out a much wider range of functions including classification tasks with comparable performance to the state-of-the-art DNABERT-2 and HyenaDNA, as well as regression and generation tasks that are not accomplishable by these tools.

## Methods and applications

Our GenomicLLM model was trained using three different datasets: GenomicLLM_GRCh38, Genome Understanding Evaluation (GUE) and GenomicBenchmarks (HyenaDNA) (Grešová, et al., 2023). After data processing and tokenization, our model was trained to perform a variety of tasks for genomic sequence analysis (Supplementary Figure 1 and Supplementary table 1). A detailed description of data processing, tokenization and model training can be found in the Supplementary material.

## Datasets

### GenomicLLM_GRCh38

A compilation of sequence and annotation information based on the NCBI Genome assembly GRCh38. The dataset comprises five subsets, namely Human Splice Site, Human Enhancers NCBI, Human ORF Dataset, Human Genetic Code Translation Dataset, and Human Gene Annotation And Sequence.

### Genome Understanding Evaluation

A dataset from DNABERT-2 (Zhou, et al., 2023), and we utilized three subsets: Promoter detection (Human), Core promoter detection (Human) and Transcription factor binding site prediction (Human).

### GenomicBenchmarks (HyenaDNA)

A dataset intended for genomic sequence classification. We utilized five subsets: Human Enhancers Cohn, Human Enhancers Ensembl, Human Regulatory, Human Nontata Promoters and Human OCR Ensembl.

## Tasks

### Classification tasks

**Human Splice Site Prediction** aims to predict the presence of splice sites in the human genome, thereby understanding how different transcripts can be produced through splicing and assisting in identifications of disease-causing mutations and the underlying mechanisms. For this task, we used the Human Splice Site dataset from GenomicLLM_GRCh38, which consists of 400-bp sequences extracted from the human reference genome. The dataset comprises three classes: donor site, acceptor site, and non-splice site. Donor site and acceptor site data are centered around splice sites, extending 200 bp upstream and downstream, while non-splice site data does not include splice sites.

**Human Enhancers Detection** allows binary classification to distinguish between enhancer sequences and non-enhancer sequences. For this task, we utilized three datasets: Human Enhancers NCBI from GenomicLLM_GRCh38, and Human Enhancers Cohn and Human Enhancers Ensembl from GenomicBenchmarks. In the Human Enhancers NCBI dataset, positive samples are sequences annotated as “Enhancer” in the GRCh38 annotation information, while negative samples are randomly extracted sequences from the rest of the GRCh38 reference genome. These negative samples match the length distribution of positive samples and do not overlap with positive samples.

**Human Promoters Detection** allows binary classification to distinguish between promoter sequences and non-promoter sequences. For this task, we utilized three datasets: Human Nontata Promoters from GenomicBenchmarks, and Promoter detection (Human) and Core promoter detection (Human) from GUE. Each sub-dataset contains slightly different promoter data. The Human Nontata Promoters dataset comprises promoters without the TATA-box, with sequences ranging from 200bp upstream to 50bp downstream of the transcription start site, totaling a length of 251bp. In the Promoter detection (Human) dataset, promoters span from 249bp upstream to 50bp downstream of the transcription start site, totaling 300bp in length. The Core promoter detection (Human) dataset includes promoters spanning from 34bp upstream to 35bp downstream of the transcription start site, totaling 70bp in length.

**Human OCR Detection** allows binary classification to distinguish between Open Chromatin Region (OCR) sequences and non-OCR sequences. The dataset utilized for this task is Human OCR Ensembl from GenomicBenchmarks.

**Human Regulatory Detection** allows ternary classification to distinguish between Enhancer sequences, Promoter sequences, and OCR sequences. The dataset used for this task is the Human Regulatory dataset from GenomicBenchmarks.

**Human Transcription Factor Binding Site Prediction** allows binary classification to predict whether a sequence is a Transcription Factor Binding Site. The dataset used for this task is the Transcription factor binding site prediction (Human) from GUE.

**Human Open Reading Frame (ORF) Detection** is to identify which of the three reading frames within a nucleotide sequence yields the functional protein. The dataset used for this task is the Human ORF Dataset from GenomicLLM_GRCh38.

**Human Seq Direction Detection** allows binary classification to identify whether a sequence is in the forward (5’ to 3’) or reverse (3’ to 5’) direction. The dataset used for this task consists of randomly extracted sequences combined with textual information from Human Gene Annotation And Sequence under GenomicLLM_GRCh38.

**Human Gene Biotype Detection** allows binary classification to identify whether a sequence belongs to a protein-coding gene or a non-coding gene (including pseudogenes, lncRNAs, miRNAs, etc.). The dataset utilized for this task comprises randomly extracted sequences combined with textual information from Human Gene Annotation And Sequence under GenomicLLM_GRCh38.

### Regression task

**Guanine-cytosine Content Prediction** aims to calculate the GC content of the input sequence. This task uses sequences randomly extracted from the Human Gene Annotation And Sequence under GenomicLLM_GRCh38, combined with textual information.

### Generation tasks

**Generation of Amino Acid Sequence from Nucleotide Sequence (NT2AA)** generates the corresponding amino acid sequences based on the nucleotide sequences. The dataset used for this task is the Human Genetic Code Translation Dataset from GenomicLLM_GRCh38, comprising CDS sequences extracted from protein-coding genes in the human reference genome along with their corresponding protein sequences.

**Human Reverse Complement Sequence Generation** generates the reverse sequence, the complementary sequence, or the reverse complementary sequence based on the input sequence. The dataset used for this task comprises randomly extracted sequences from Human Gene Annotation And Sequence under GenomicLLM_GRCh38.

**Human Gene Description Generation** generates descriptive information about a gene based on a combination of partial gene sequences and accompanying annotations like IDs. The dataset used for this task comprises randomly extracted sequences combined with textual information from Human Gene Annotation And Sequence under GenomicLLM_GRCh38.

### Reading comprehension task

**Textual Sequence Reading Comprehension** aims to enable the model to comprehend both textual and sequence-based information within the input and extract the correct answer for the input question. The dataset used for this task comprises randomly extracted sequences combined with textual information from Human Gene Annotation And Sequence under GenomicLLM_GRCh38.

## Results and discussion

In our GenomicLLM tool, the input consists of a mixed corpus that starts with either a DNA/protein sequence or gene/sequence annotation information, followed by a specific request, and ends with the “=” sign, while the output returns the answer to the input request, similar to a Q&A format. Examples of the inputs and outputs for each tasks can be found in Supplementary table S1.

GenomicLLM can perform a broader variety of tasks than DNABERT-2 and HyenaDNA with comparable performance in classification tasks. GenomicLLM outperforms both DNABERT-2 and HyenaDNA in 7 out of 13 classification tasks on the GUE and the GenomicBenchmarks datasets (Table 2). Moreover, GenomicLLM can also perform more tasks that cannot be done in DNABERT-2 or HyenaDNA, including more classification and regression tasks like identifications of the correct ORFs, forward vs reverse strands and protein-coding vs non-protein-coding sequences, GC content calculation (Table 3), as well as generation tasks like nucleotide sequence-to-amino acid sequence translation, reverse complement sequence generation, reading comprehension and gene information generation (Table 4) with outstanding performance.

**Table 2:**
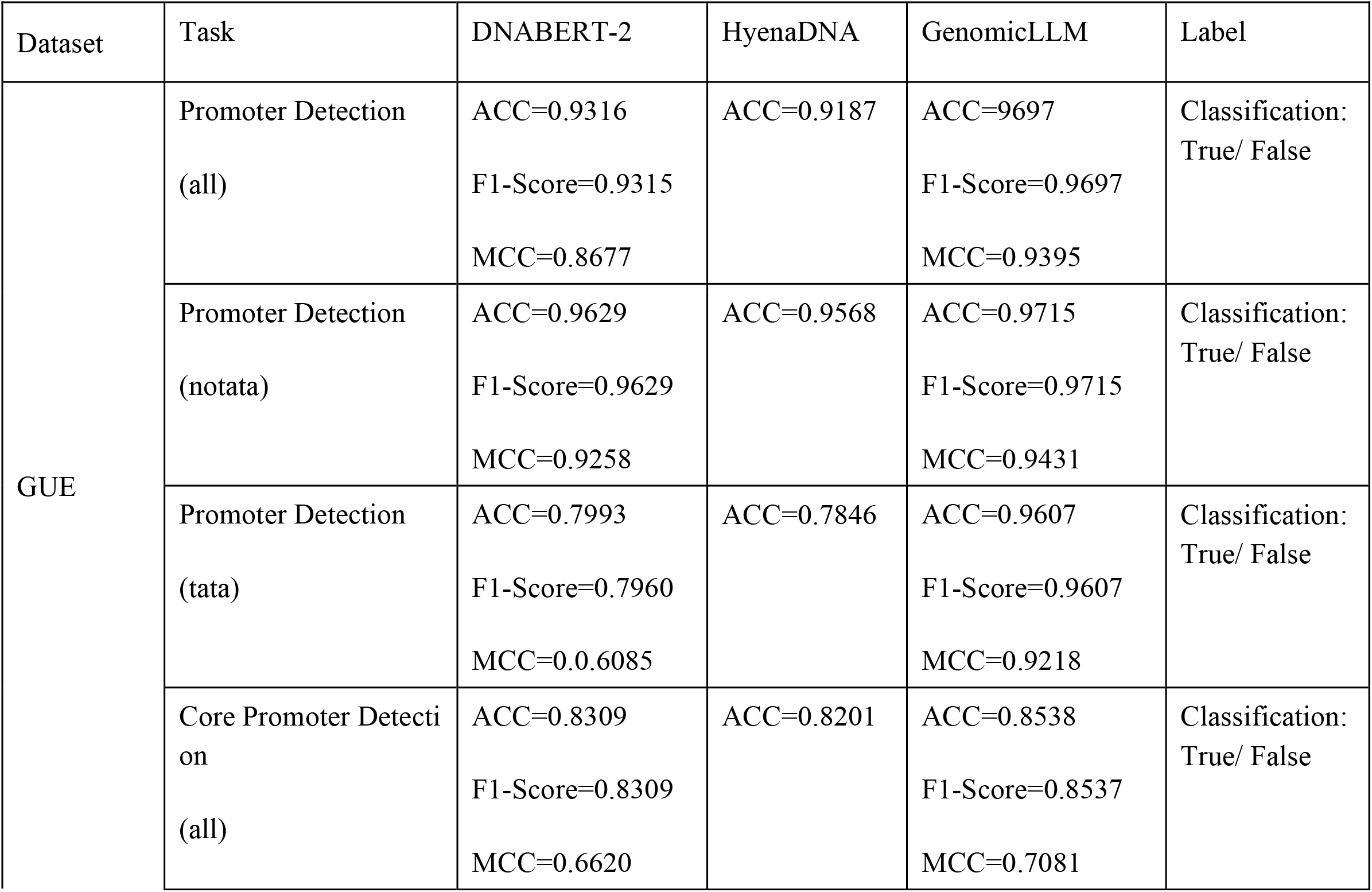
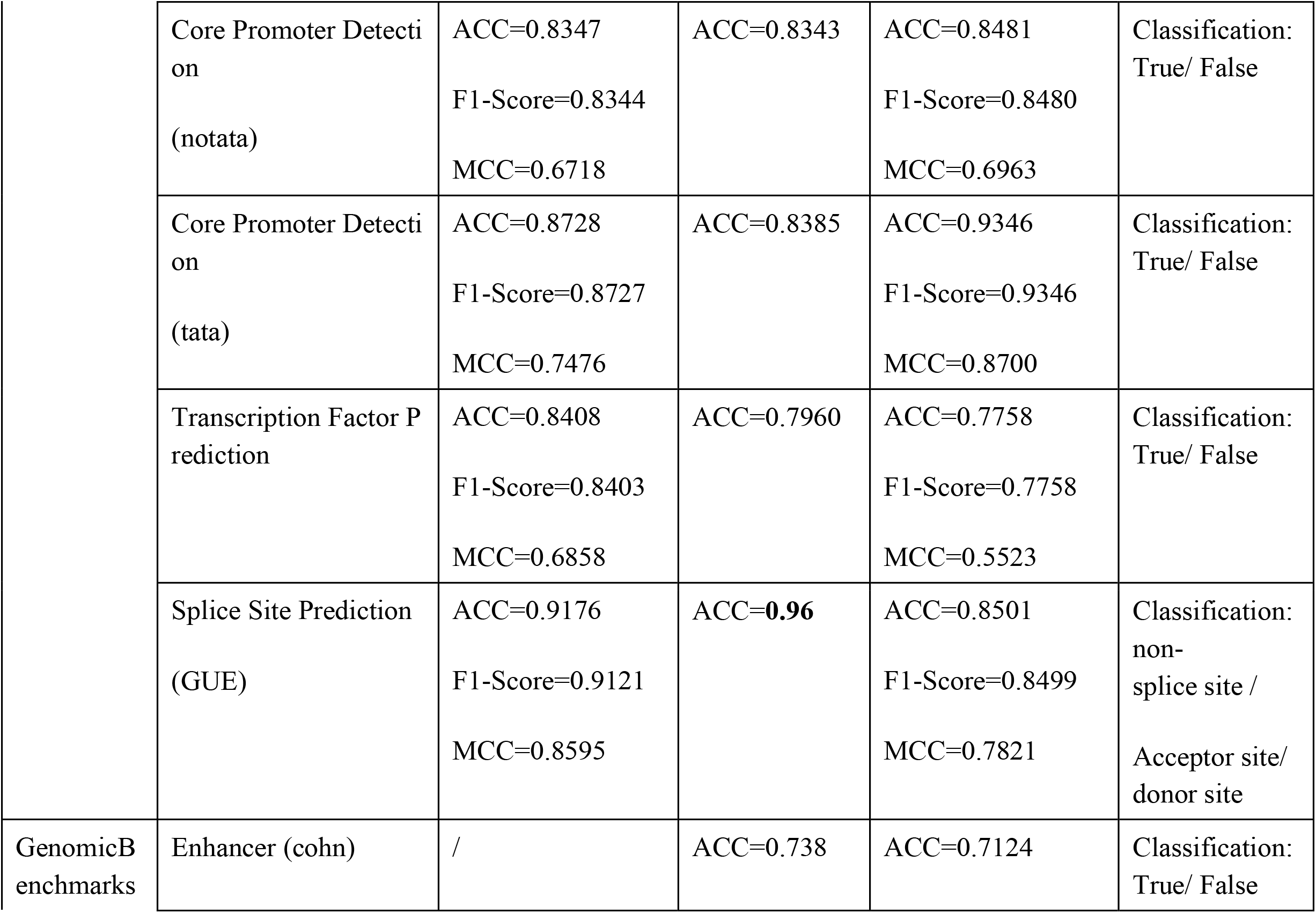
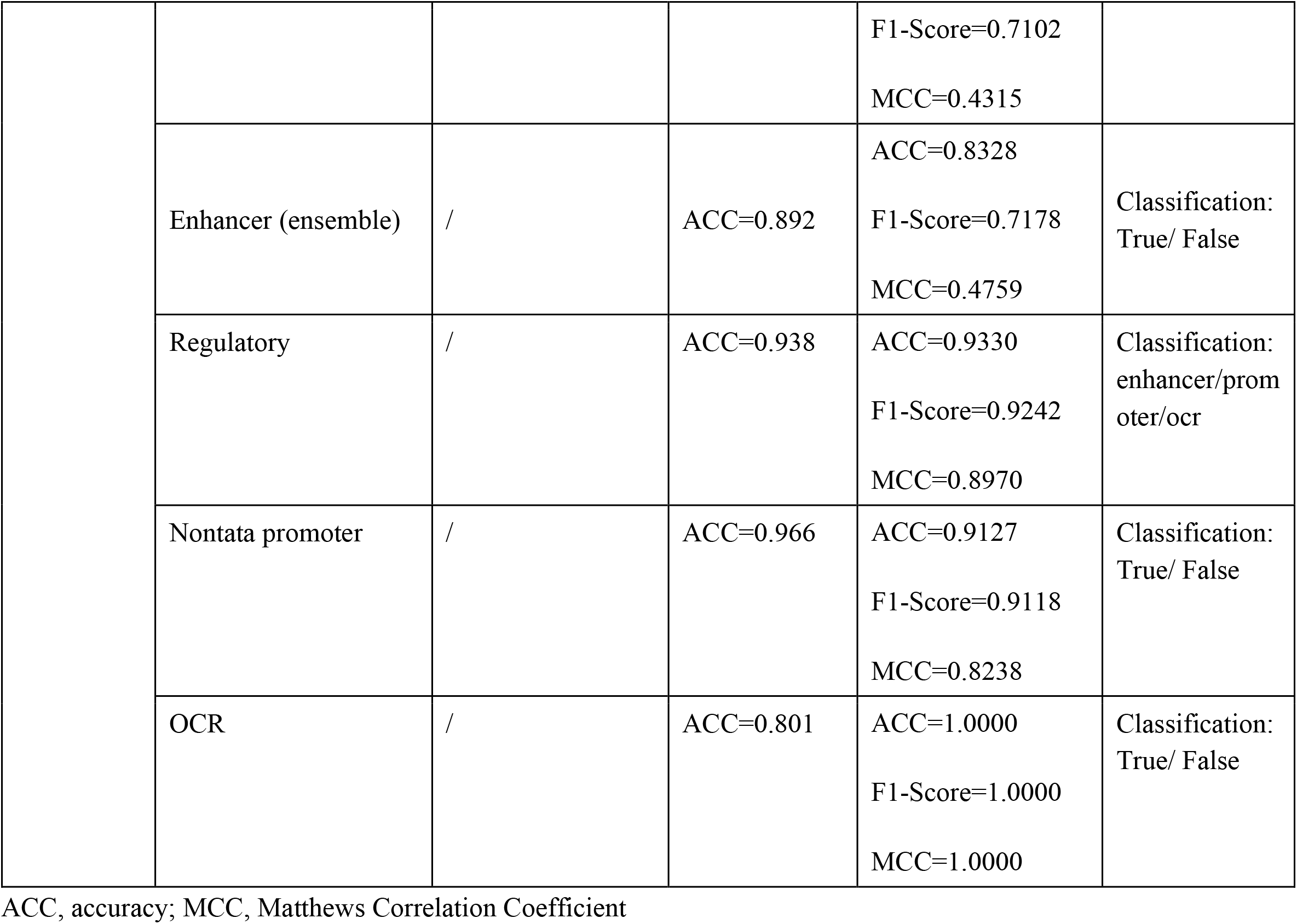
Comparisons between GenomicLLM, DNABERT-2 and HyenaDNA in classification tasks on the GUE and the GenomicBenchmarks datasets.

**Table 3:** The performance of GenomicLLM in classification and regression tasks on the GenomicLLM_GRCh38 dataset.

| Dataset | Task | GenomicLLM | Label |
| --- | --- | --- | --- |
| GenomicLLM_<br>GRCh38 | Splice Site Prediction<br>(GenomicLLM) | ACC=0.9844 F1-Score=0.9845 MCC=0.9766 | Classification: True/ False |
|  | Splice Site Prediction<br>(GenomicLLM+GUE) | ACC=0.9660 F1-Score=0.9665 MCC=0.9490 | Classification: True/ False |
|  | enhancer<br>(GenomicLLM) | ACC=0.8798 F1-Score=0.8791 MCC=0.7639 | Classification: True/ False |
|  | enhancer<br>(GenomicLLM+GenomicBenchmarks) | ACC=0.8287 F1-Score=0.8157 MCC=0.6446 | Classification: True/ False |
|  | ORF | ACC=0.9997 F1-Score=0.9997 MCC=0.9995 | Classification: True/ False |
|  | sequence direction | ACC=1.0000 F1-Score=1.0000 MCC=1.0000 | Classification: True/ False |
|  | gene biotype | ACC=0.7566 F1-Score=0.6852 MCC=0.4424 | Classification: True/ False |
|  | GC content | Pearson R = 1.0000<br>MAE = 0.0010 | Regression |

ACC, accuracy; MCC, Matthews Correlation Coefficient

**Table 4:** The performance of GenomicLLM in generation tasks on the GenomicLLM_GRCh38 dataset.

| Task | Data size | ACC | BLEU-1 | BLEU-2 | BLEU-3 | BLEU-4 | ROUGE<br>_1 | ROUGE<br>_2 | ROUGE_<br>L |
| --- | --- | --- | --- | --- | --- | --- | --- | --- | --- |
| reading comprehension | 79 | 1.0000 | 1.0000 | 1.0000 | 1.0000 | 1.0000 | 1.0000 | 1.0000 | 1.0000 |
| annotation:description | 100 | 0.5100 | 0.6853 | 0.6255 | 0.6040 | 0.5878 | 0.7165 | 0.6345 | 0.7053 |
| protein seq | 1716 | 0.9930 | 0.9999 | 0.9998 | 0.9998 | 0.9997 | 1.0000 | 0.9999 | 1.0000 |
| reverse sequence | 4309 | 1.0000 | 1.0000 | 1.0000 | 1.0000 | 1.0000 | 1.0000 | 1.0000 | 1.0000 |
| complementary sequence | 4250 | 0.9993 | 0.9999 | 0.9999 | 0.9999 | 0.9999 | 1.0000 | 1.0000 | 1.0000 |
| reverse complementary sequence | 4382 | 1.0000 | 1.0000 | 1.0000 | 1.0000 | 1.0000 | 1.0000 | 1.0000 | 1.0000 |

ACC, accuracy; BLEU, Bilingual Evaluation Understudy (Papineni, et al., 2002); ROUGE, Recall Oriented Understudy for Gisting Evaluation (Lin, 2004)

Our GenomicLLM model demonstrates a successful large language model with a mixture of gene sequences and natural language corpus, enabling a wider range of applications compared to models that have undergone downstream task fine-tuning, and comparable performance to the state-of-the-art tools. Genomic LLM, as a pre-trained model for mixed corpora, is more convenient to expand and has better application prospects than pure gene sequence models like DNABERT-2.

SentencePiece is a language-agnostic tokenizer used to segment text data into subwords or character sequences (Kudo and Richardson, 2018). It learns a fixed-size, variable-length vocabulary of tokens based on the co-occurrence frequency of characters. It can be employed for various natural language processing tasks. We first attempted to tokenize DNA sequences and text using SentencePiece with Byte Pair Encoding (BPE) only. However, due to the intrinsic differences, such as in letter frequencies and meaning, between gene sequences and natural language texts, we found that this method would create arbitrary k-mer words in DNA sequences, which can actually make some tasks, such as gene sequence generation and translating nucleotides into amino acids, very difficult. Hereby, we designed a hybrid tokenization approach that adopts different processing methods for different corpora to better extract information from mixed corpora.

In order to reduce the difficulty of understanding natural language texts and the cost of model training from scratch, we have defined a relatively simple grammar, similar to symbolic languages. Currently, only natural language dialogues with specific genomic terms are supported due to the use of a nano-LLaMA2 network. In the future, we plan to add genomic sequence and non-sequence corpora and perform fine-tuning based on a larger natural language model, to create an all-in-one genomic sequence analysis tool that can understand natural language questions and answer in a daily-conversation style.

## Competing interests

The authors declare that they have no competing interests.

## Authors’ contributions

All the authors contribute to designing the product, analyzing data and writing the manuscript, Huaqing Liu, Shuxian Zhou and Peiyi Chen contribute to AI model building, Jiahui Liu contributes to dataset processing and bioinformatic analysis.

## Acknowledgements

These should be included at the end of the text and not in footnotes. Please ensure you acknowledge all sources of funding, see funding section below.

Details of all funding sources for the work in question should be given in a separate section entitled ‘Funding’. This should appear first in the ‘Acknowledgements’ section.

## Funding

This work was supported by Cyagen Ltd.

## Notes

### Competing Interest Statement

The authors have declared no competing interest.

