## Supplementary material for "Exploring Genomic Large Language Models: Bridging the Gap between Natural Language and Gene Sequences"

### Tokenization

We used the SentencePiece tokenizer (Kudo and Richardson, 2018) and a hybrid tokenization approach that adopts different processing methods for different corpora. We first performed predefined tokens in the vocabulary, including genome specific nouns, numbers, symbols, kmer (1<=k<=3), three-letter amino acid abbreviations, etc., then, using the sense piece method, we automatically fit on the training set to obtain a mixed-type token.

### Data processing

Due to imbalanced data, to better train the model and avoid the model focusing on learning certain tasks, we assigned different weights to different datasets to achieve the effect of balancing the data. The calculation method for weight is as follows:

$$w_i = \frac{N_{all}}{N_i}$$

Where $w_i$ is the sampling weight of the i-th dataset, $N_{all}$is the total amount of data in the dataset, and $N_i$is the amount of data in the i-th dataset.

The xx data is composed of various gene descriptions and information combinations. In order to improve the diversity of the corpus, the xx data were processed as follows: randomly select a type of description or information as a problem, and randomly select several types as the associated text of the problem to form a new piece of data. Using this method, real-time data are generated during model training. Considering the cost of model training and ensuring the effectiveness of the corpus, the length of gene sequences in this experiment corpus should not exceed 512 tokens and not be less than 4 tokens, and the natural language text should not exceed 256 tokens and not be less than

2 tokens. For duplicate parts in different datasets, we retained the data in GUE and HyenaDNA and deleted the data in xx dataset.

#### Model training and inference

To quickly validate our ideas without using too much computing resources, we first used a nano-LLaMA2 network with only 88M parameters instead of the complete LLaMA-7B network.

**Model parameters**

| Network | nano-LLaMA2 |
|---|---|
| Number of transformer block | 12 |
| Total dimension of attention | 768 |
| Number of attention heads | 12 |
| Max input tokens | 512 |
| Model parameters | 88M |

We trained the model by Next Token Prediction task with cross-entropy loss. We used a simple token "=" as the separator between the question and the answer. Before calculating the total loss value, tokens were splited by the last '=' into two groups, namely the question and the answer. The average loss of answer tokens were given a weight of 1.0, and the average loss of question tokens 0.2. The weighted sum is calculated as the total loss value. When adding "=" at the end of the input during inference, the model will generate answers through autoregression until 'eos' (end of sentence) is generated or the length of the generated text reaches the maximum length limit. We trained the model for 500,000 steps using the AdamW optimizer (Loshchilov and Hutter, 2017) on a NVIDIA A100 GPU. The learning rate increased linearly from 0 to 5e-4 for the first 3000 steps and decreased following a cosine decay for the rest steps.

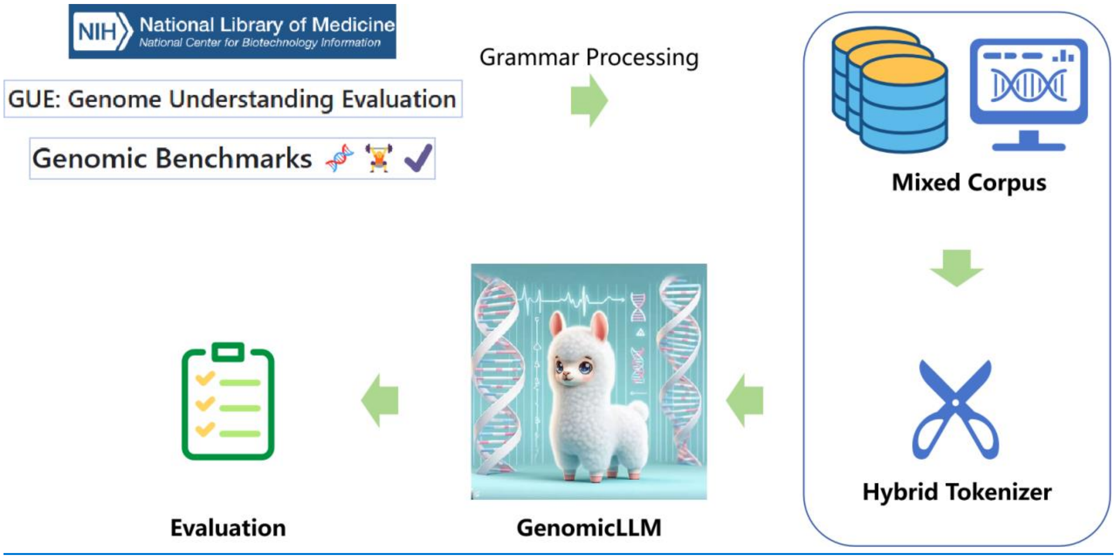


**Figure S1.** Overview of GenomicLLM training

**Table S1.** Overview of the Datasets and tasks.

| Datasets | Tasks | Training | Validation | Test |
|---|---|---|---|---|
| GUE | Promoter Detection (all) | 47356 | 5920 | 5920 |
| GUE | Promoter Detection (notata) | 42542 | 5307 | 5307 |
| GUE | Promoter Detection (tata) | 4904 | 613 | 613 |
| GUE | Core Promoter Detection (all) | 47356 | 5920 | 5920 |
| GUE | Core Promoter Detection (notata) | 42542 | 5307 | 5307 |
| GUE | Core Promoter Detection (tata) | 4904 | 613 | 613 |
| GUE | Transcription Factor Prediction | 128344 | 4600 | 4906 |
| GUE | Splice Site Prediction (GUE) | 35717 | 4354 | 4324 |
| GenomicBenchmarks | enhancer | 97017 | 24255 | 24693 |
| GenomicBenchmarks | regulatory | 185072 | 46268 | 57713 |
| GenomicBenchmarks | Nontata promoter | 21677 | 5420 | 9034 |

| Datasets | Tasks | Training | Validation | Test |
|---|---|---|---|---|
| GenomicBenchmarks | OCR | 111792 | 27949 | 34941 |
| GenomicLLM_GRCh38 | SSP | 650664 | 36148 | 36148 |
| GenomicLLM_GRCh38 | enhancer | 117584 | 14698 | 14696 |
| GenomicLLM_GRCh38 | ORF | 79127 | 9891 | 9891 |
| GenomicLLM_GRCh38 | aa2nt | 60090 | 3338 | 3339 |
| GenomicLLM_GRCh38 | reverse sequence | random | random | random |
| GenomicLLM_GRCh38 | complementary sequence | random | random | random |
| GenomicLLM_GRCh38 | reverse complementary sequence | random | random | random |

| Datasets | Tasks | Training | Validation | Test |
|---|---|---|---|---|
| GenomicLLM_GRCh38 | direction | random | random | random |
| GenomicLLM_GRCh38 | Gene biotype | 12220 | 1528 | 1528 |
| GenomicLLM_GRCh38 | GC content | random | random | random |
| GenomicLLM_GRCh38 | reading comprehension | random | random | random |
| GenomicLLM_GRCh38 | description | random | random | random |
